# Comparing the differentiation potential of *Brachyury*^+^ mesodermal cells generated from 3-D and 2-D culture systems

**DOI:** 10.1101/239913

**Authors:** Jing Zhou, Antonius Plagge, Patricia Murray

**Affiliations:** Institute of Translational Medicine, University of Liverpool, Liverpool L69 3BX, UK

## Abstract

Mesodermal populations can be generated *in vitro* from mouse embryonic stem cells (mESCs) using three-dimensional (3-D) aggregates called embryoid bodies or two-dimensional (2-D) monolayer culture systems. Here, we investigated whether *Brachyury*-expressing mesodermal cells generated using 3-D or 2-D culture systems are equivalent, or instead, have different properties. Using a *Brachyury*-GFP/E2-Crimson reporter mESC line, we isolated *Brachyury-*GFP^+^ mesoderm cells using flow-activated cell sorting and compared their gene expression profiles and *ex vivo* differentiation patterns. Quantitative RT-PCR analysis showed significant up-regulation of *Cdx2*, *Foxf1* and *Hoxb1* in the *Brachyury*-GFP^+^ cells isolated from the 3-D system compared with those isolated from the 2-D system. Furthermore, using an e*x vivo* mouse kidney rudiment assay, we found that irrespective of their source, *Brachyury*-GFP^+^ cells failed to integrate into developing nephrons, which are derived from the intermediate mesoderm. However, *Brachyury*-GFP^+^ cells isolated under 3-D conditions appeared to differentiate into endothelial-like cells within the kidney rudiments, whereas the *Brachyury*-GFP^+^ isolated from the 2-D conditions only did so to a limited degree. The high expression of *Foxf1* in the 3-D *Brachyury*-GFP^+^ cells combined with their tendency to differentiate into endothelial-like cells suggests these mesodermal cells may represent lateral plate mesoderm.

## 1. Introduction

The formation of the primitive streak (PS) marks the onset of antero-posterior axis determination in the developing mouse embryo (Stern, 2004; Rodriguez *et al*., 2005). The epiblast cells egress through the PS to generate the nascent mesoderm in-between the primitive ectoderm and the overlying visceral endoderm. *Brachyury* (*Bra*, also known as *T*) is the key marker of the entire PS and is a pan mesodermal marker that is expressed in the posterior epiblast, PS, node, notochord, allantois and tail bud (Wilkinson *et al*., 1990; Kispert and Herrmann, 1994; Conlon *et al*., 1995; Kispert *et al*., 1995; King *et al*., 1998; Showell *et al*., 2004; Papaioannou, 2014; Concepcion and Papaioannou, 2014).

Following gastrulation, the *Bra*^+^ nascent mesoderm generates (i) paraxial mesoderm, which gives rise to the somites; (ii) lateral plate mesoderm, which gives rise to the heart, vessels, haematopoietic stem cells and endothelial cells; and (iii) intermediate mesoderm, which gives rise to the urogenital system (Gilbert, 2010; Wolpert *et al*., 2015). The intermediate mesoderm then becomes further specified to anterior intermediate mesoderm that gives rise to the ureteric bud (UB), and posterior intermediate mesoderm that gives rise to the metanephric mesenchyme (MM) (Little *et al*., 2016). The UB and MM generate the collecting ducts and nephrons, respectively, of the mature kidney (Pietilä and Vainio, 2014; Little *et al*., 2016).

The small size and inaccessibility of the peri-implantation mouse embryo makes it difficult to study. However, the isolation of embryonic stem cells (ESCs) from mouse blastocysts in the 1980s (Evans and Kaufman, 1981; Martin, 1981) has provided an alternative model for studying the early development of the mouse embryo.

When cultured in suspension, mESCs spontaneously form spheroid multicellular aggregates called embryoid bodies (EBs) (Wobus *et al*., 1984; Doetschman *et al*., 1985; Robertson, 1987; Murray and Edgar, 2004). A typical EB has an outer layer of primitive endoderm, an inner layer of primitive ectoderm, a basement membrane separating them, as well as a central cavity that resembles the proamniotic cavity (Shen and Leder, 1992). The primitive ectoderm differentiates to generate derivatives of definitive ectoderm, endoderm and mesoderm (Wobus *et al*., 1984; Doetschman *et al*., 1985; Keller *et al*., 1993). Therefore, EBs can recapitulate some aspects of peri-implantation mouse development and provide an excellent model system for studying these early events (Wobus *et al*., 1984; Doetschman *et al*., 1985; Robertson, 1987).

However, the heterogeneous nature of the EBs means that the extent of differentiation towards any specific cell type can vary considerably depending on culture conditions, and can even vary between EBs cultured under the same culture conditions. The complex 3-D structure also hinders the visualisation of the differentiation process at an individual cell level. For this reason, various 2-D differentiation protocols have been developed to direct differentiation to specific cell-types more efficiently. Several studies have demonstrated *in vitro* derivation of monolayer mESCs into lineages of neural progenitors, endothelial cells, osteochondrogenic and myogenic cells using chemically defined media (Ying and Smith, 2003; Sakurai *et al*., 2009; Blancas *et al*., 2011; Blancas *et al*., 2013). Recently, Turner *et al* showed that *Activin/Nodal* and *Wnt* signalling pathways promote mesoderm formation in monolayer mESC culture, with the mesodermal cells differentiated from mESCs displaying *Bra* expression, similarly to the nascent mesoderm that develops in the primitive streak of developing mouse embryos and of ‘gastrulating’ EBs. By using a combination of Activin A (*Activin/Nodal* agonist) and Chiron (*Wnt3a* agonist), this group developed a highly efficient strategy for inducing E14 mESCs to differentiate into nascent mesoderm. After 2-day culture in neural differentiation medium and a further 2-day culture in medium supplemented with Activin A and Chiron, robust *Bra* expression was observed in over 90% of the population (David Turner, University of Cambridge, personal communication) (Turner *et al*., 2014a,b).

Although mesoderm differentiation occurs within both the 3-D EB and 2-D mESC culture systems, it is not clear whether the differentiated cells (e.g. mesodermal cells) that are generated by the 2-D protocols are equivalent to those that form in EBs. In the mouse embryo, the fate of the *Bra*^+^ cells is determined by the microenvironment that the cells find themselves in following their migration from the primitive streak (Gilbert, 2010). This cannot be replicated using *in vitro* culture systems, which raises the question of whether the *Bra*^+^ cells generated *in vitro* are equivalent to nascent mesoderm, or instead, are partially committed to a specific mesodermal lineage. For instance, the Little group have previously reported that BRA^+^ cells derived from human ESCs have a tendency to spontaneously differentiate into FOXF1^+^ lateral plate mesoderm when cultured in the absence of exogenous growth factors (Takasato *et al*., 2014). This observation highlights the fact that the differentiation potential of Bra+ cells generated in vitro is likely to be influenced by the specific culture conditions used. We have previously shown that *Bra*^+^ mesodermal cells isolated from mESC-derived EBs were able to integrate into the developing UB and MM of mouse kidney rudiments and generate specialised renal cells (Rak-Raszewska *et al*., 2012). However, in this previous study, the EBs from which the *Bra*^+^ mesodermal cells were isolated did not mimic early embryo development, in that they did not form a primitive ectoderm epithelium, nor a proamniotic cavity. In the present study, we aimed to investigate whether *Bra*^+^ cells generated using the recently described 2-D culture system, and those derived from cavitating EBs, express similar lineage-specific genes, and have similar developmental potential to those derived from non-cavitating EBs. In order to do this, we have generated a *Bra-GFP/Rosa26-E2C* mESC reporter line (Zhou *et al*., in press) that will allow us to isolate the GFP-expressing nascent mesodermal cells from both systems so that their gene expression can be analysed using RT-PCR and their developmental potential can be assessed by investigating their fate following incorporation into mouse kidney rudiments *ex vivo* (Unbekandt and Davies, 2010; Kuzma-Kuzniarska *et al*., 2012; Rak-Raszewska *et al*., 2012; Ranghini *et al*., 2013; Dauleh *et al*., 2016).

## 2. Results

### 2.1 Mesoderm development within EBs is affected by seeding density

The *Bra-GFP/Rosa26-E2C* mESCs were plated at different densities and cultivated for 7 days in EB medium. At densities of 2.5×10^5^ and 1.25×10^5^ cells mL^−1^, cavitated EBs could be observed by day 4, but at the lower seeding density of 6.25×10^4^ cells mL^−1^, most EBs failed to cavitate, even by day 7 (Fig. 1). Mesoderm development was identified in all conditions by GFP fluorescence, but the expression patterns were different. At 6.25×10^4^ cells mL^−1^, GFP was expressed at an earlier stage and peaked on day 4 before decreasing. In contrast, at higher densities, GFP became visible at day 4 or later and the fluorescence signal increased from day 4 to 7, but there appeared to be more GFP^+^ cells in the 1.25×10^5^ cells mL^−1^ EBs (Fig. 1). Therefore, given that the EBs developing in the 1.25×10^5^ cells mL^−1^ density cultures appeared to be typical cavitating EBs that contained a high proportion of GFP^+^ cells, we used this plating density in all future experiments. To investigate if E2C expression affected mesoderm differentiation, immunostaining of *Bra-GFP/Rosa26-E2C* EB sections was performed to confirm that the GFP^+^ cells within the EB expressed E2C. The results showed that all cells within the *Bra-GFP/Rosa26-E2C* EBs continued to express E2C, including the GFP^+^ mesodermal cells, indicating that E2C expression did not inhibit mesoderm differentiation (Fig. 1).

**Fig. 1.**
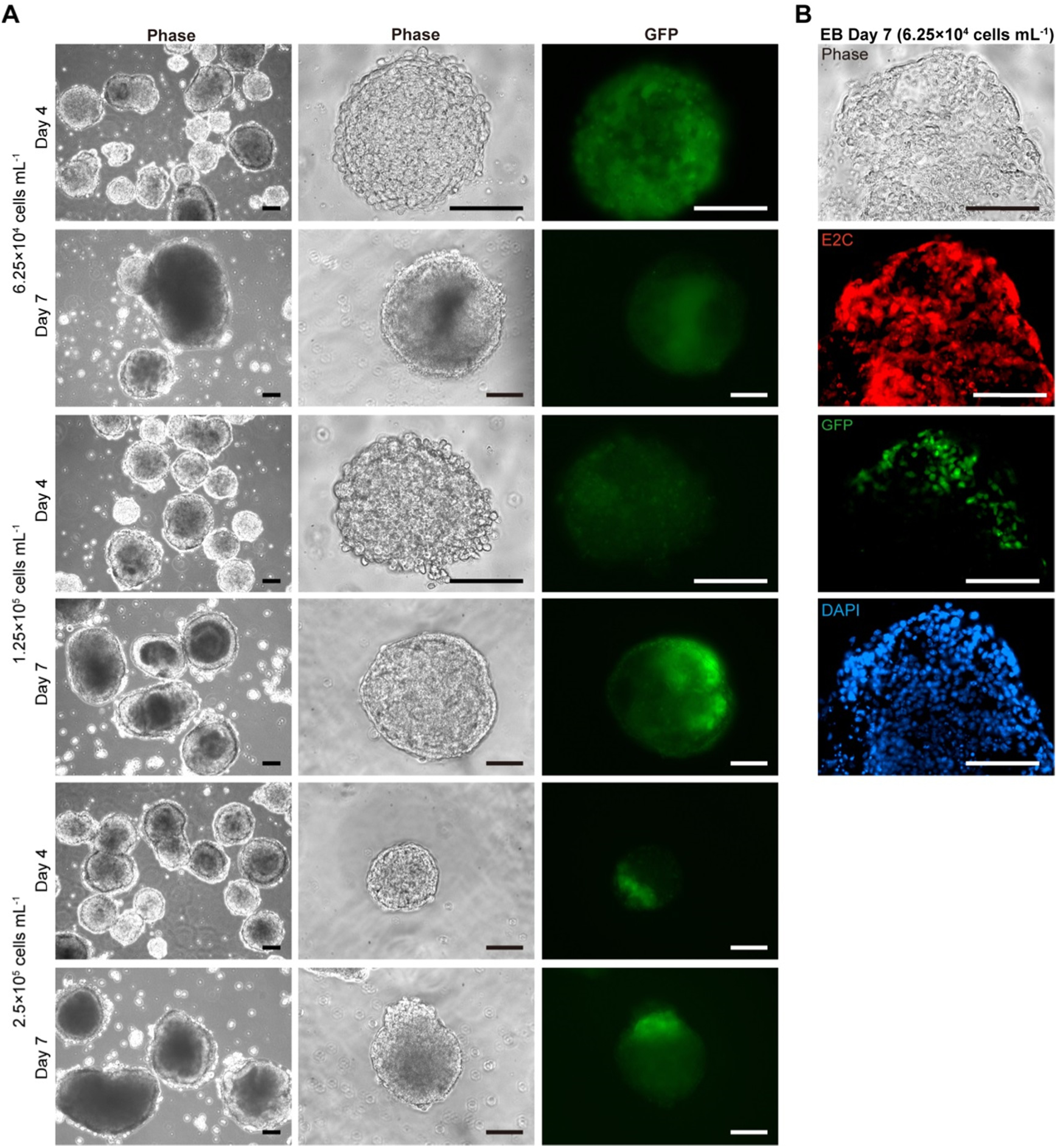
Representative fluorescence and phase contrast photomicrographs of mesoderm development within EBs derived from *Bra-GFP/Rosa26-E2C* mESCs at different seeding densities cultured for up to 7 days. (A) EB morphology was examined on days 4 and 7. The majority of EBs derived from mESCs plated at densities of 2.5×10^5^ and 1.25×10^5^ cells mL^−1^ showed evidence of cavitation, whereas cavitated EBs were less abundant in the lower density culture (6.25×10^4^ cells mL^−1^). Maximal levels of GFP expression were observed in day 7 EBs derived from the 1.25×10^5^ density cultures. (B) Immunostaining of cryo-sections of day 7 EBs for E2C, counterstained with DAPI. Representative photomicrographs of lower density culture showed that all cells within the EBs derived from the E2C-expressing mESCs stained positively for E2C (red), including the GFP^+^ (green) mesodermal cells. Data were collected from three biological replicates. Scale bars, 100 μm.

### 2.2 Comparing the timing and extent of mesodermal cell differentiation using the 3-D and 2-D culture systems

In order to accurately monitor changes in GFP expression in the developing EBs over time, *Bra-GFP/Rosa26-E2C* mESCs were plated at a density of 1.25×10^5^ cells mL^−1^ and at day 3, were embedded in a sandwich-like agarose system (2% agarose bottom layer ‒ EB ‒ 1% agarose overlay) and imaged in real-time using the Cell-IQ instrument every hour from day 3 to day 9 post plating. GFP started to be expressed on day 4 (96 h), and reached maximum levels on day 6–7. Although expression levels began to decrease at this time point, GFP^+^ cells were still present at day 9 (Fig. 2A). To quantify the proportion of mesodermal cells within the EBs, flow cytometry analysis was performed. EBs derived from the wild-type E14TG2a mESCs were used as a negative control. The results were consistent with the Cell-IQ data, and showed that the peak GFP expression was at day 6, at which time, approximately 39% of the EB population were GFP^+^ (Fig. 2B).

**Fig. 2.**
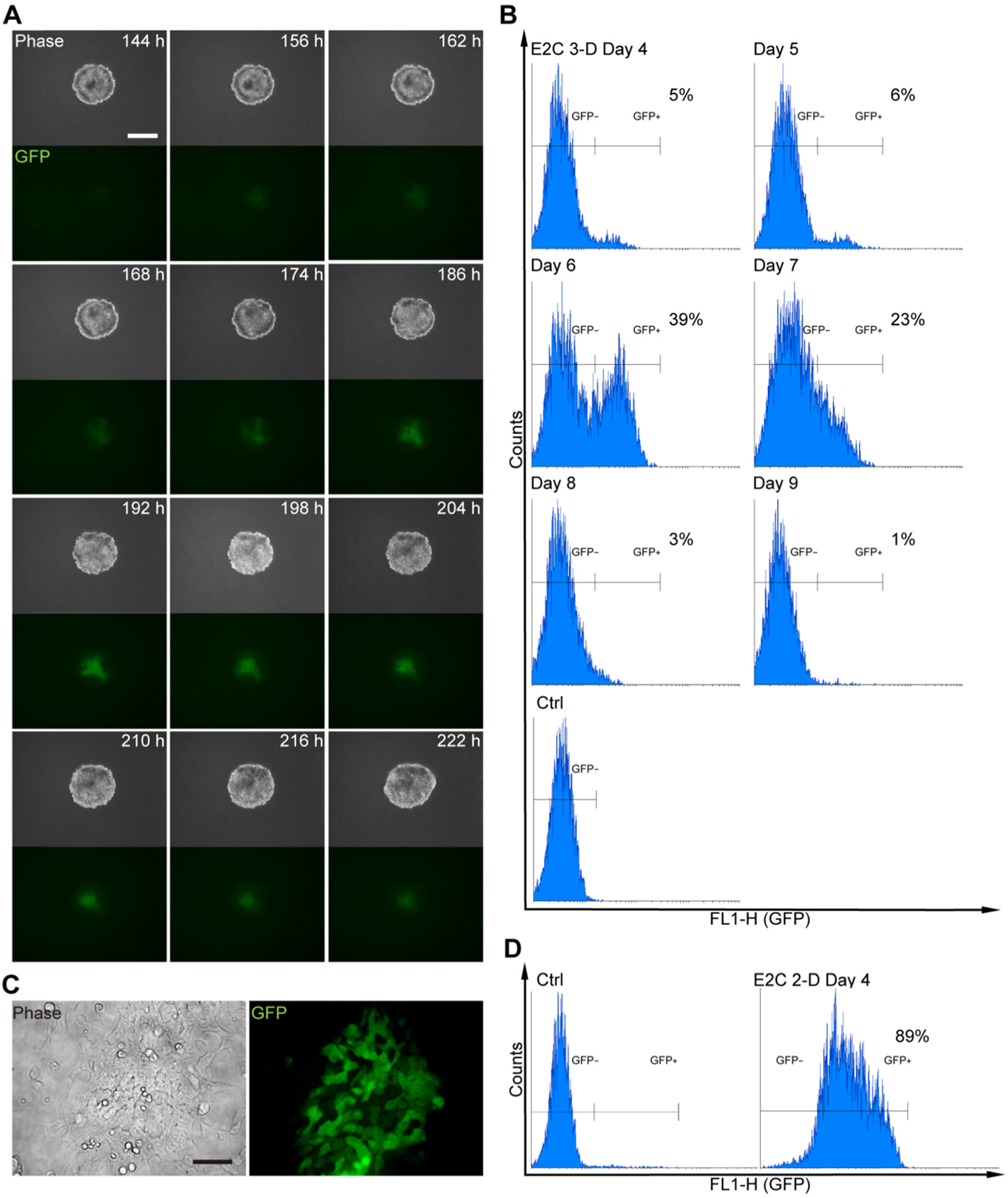
Timing and extent of GFP expression in *Bra-GFP/Rosa26-E2C* mESCs following mesodermal differentiation in 3-D and 2-D culture systems. (A) Fluorescence and phase contrast photomicrographs of EBs derived from mESCs plated at 1.25× 10^5^ cells mL^−1^. EBs were cultured for up to 9 days and imaged in real-time every hour from day 3 to day 9 post plating. GFP expression could still be detected at day 9. (B) Flow cytometry analysis of disaggregated *Bra-GFP/Rosa26-E2C* EBs at different time points revealed that GFP started to be expressed on day 4, and reached maximum levels on days 6–7. At the peak of expression (day 6), GFP^+^ cells comprised 39% of the population. (C) Representative fluorescence and phase contrast photomicrographs of mESCs following directed differentiation to mesoderm using a 2-D culture system. Four days following induction, cells no longer formed colonies, appeared differentiated, and the majority expressed GFP. (D) Flow cytometry analysis showed that ~89% of cells expressed GFP under 2-D culture conditions. Undifferentiated *Bra-GFP/Rosa26-E2C* mESCs sub-cultured in gelatinised dishes in mESC medium for 24 h prior to induction were used as a negative control. Data were collected from at least 2 biological replicates. Scale bars, 200 μm (A) and 100 μm (C).

We then determined the efficiency of the previously described 2-D culture system (Turner *et al*., 2014a,b). The *Bra-GFP/Rosa26-E2C* mESCs were cultured under differentiation conditions for 4 days, and were then screened for GFP expression. Analysis of fixed cells in culture showed that the vast majority of the population expressed GFP. Flow cytometry analysis showed that approximately 89% of the population was GFP^+^, which is consistent with the efficiency reported previously with this method (Figs 2C–D).

### 2.3 Comparing the expression profile of key genes in GFP^+^ mesodermal cells generated under 3-D and 2-D differentiation conditions

Before comparing the expression levels of the key target genes in the GFP^+^ cells isolated from the 3-D and 2-D culture systems, it was first necessary to determine the purity of the GFP^+^ cell populations isolated from each culture system. Single cell suspensions from day 6 EBs and day 4 2-D monolayer cultures were sorted by FACS and then re-analysed using the same parameters. Results showed that the proportion of GFP^+^ cells was over 94% (Fig. 3A), confirming they were pure populations.

**Fig. 3.**
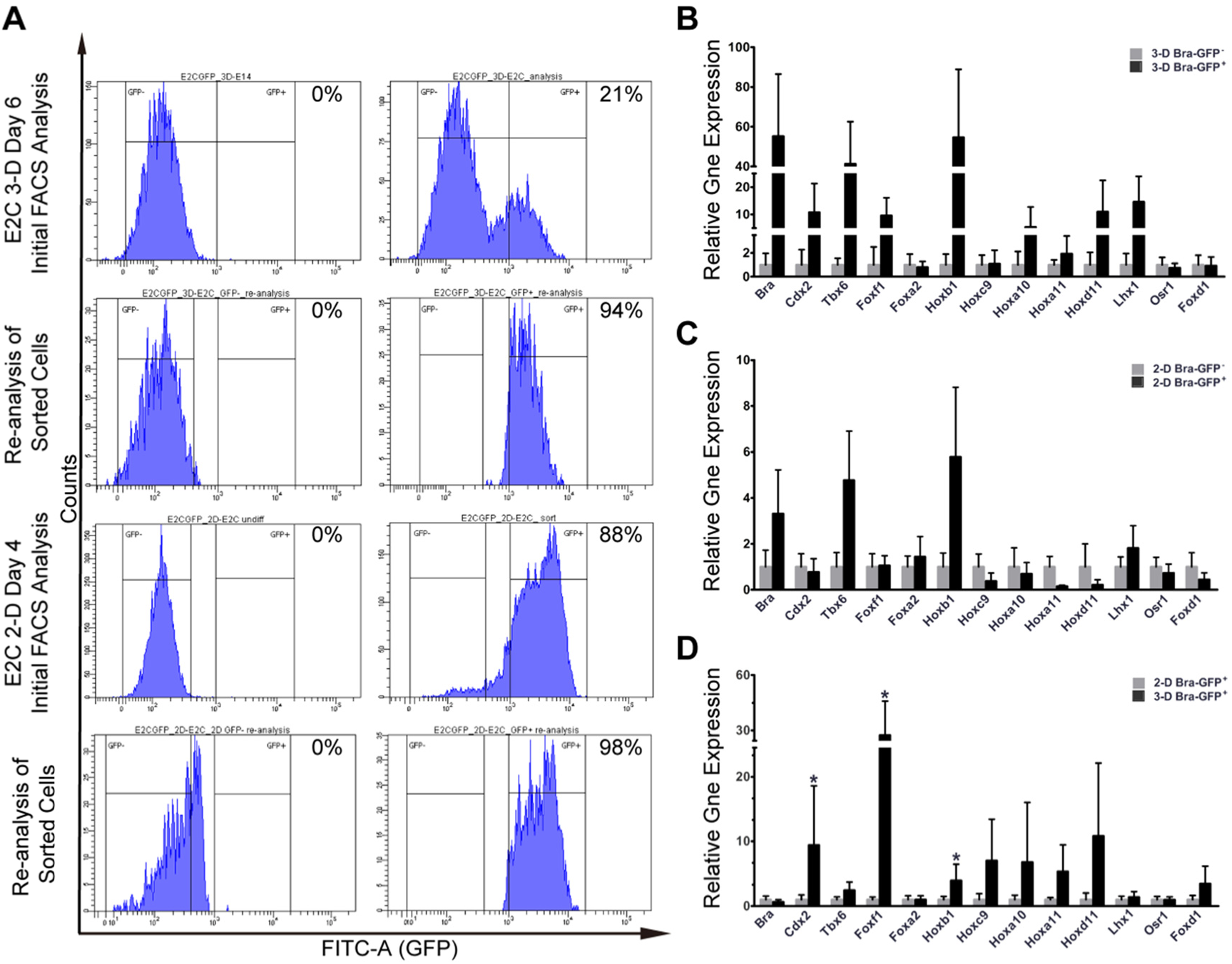
Isolation and analysis of gene expression profiles of the mesodermal and non-mesodermal populations from *Bra-GFP/Rosa26-E2C* mESCs cultured in 3-D and 2-D systems. (A) Day-6 EBs or mESCs cultured under 2D differentiating conditions for 4 days were harvested for FACS. Untransfected day-6 *E14-Bra-GFP* EBs or mESCs maintained undifferentiated in gelatinised dishes in mESC medium for 24 h prior to induction were used as negative controls. Flow cytometry was used to confirm the purity of the populations isolated using FACS. (B) Relative expression levels of mesoderm and early kidney development genes were compared between *Bra*-GFP^+^ and *Bra*-GFP^−^ populations isolated from the 3-D system (n=2 biological replicates), presented as mean±s.e.m. Data were not statistically assessed on significance due to 2 biological replicates however they gave an indication of the difference between *Bra*-GFP^+^ and *Bra*-GFP^−^ populations. (C) Relative expression levels of mesoderm and early kidney development genes were compared between *Bra*-GFP^+^ and *Bra*-GFP^−^ populations isolated from the 2-D system (n=2 biological replicates), presented as mean±s.e.m. Data were not statistically assessed on significance due to 2 biological replicates however they gave an indication of the difference between *Bra*-GFP^+^ and *Bra*-GFP^−^ populations. (D) Relative gene expression levels of mesoderm and early kidney development genes were compared between *Bra*-GFP^+^ populations isolated from 3-D system (n=3 biological replicates) and 2-D system (n=3 biological replicates), presented as mean±s.e.m. *P*<0.05 (asterisks) was considered as statistically significant (*t*-test).

In order to characterize the *Bra*-GFP+ and *Bra*-GFP− populations, quantitative real-time polymerase chain reaction (qRT-PCR) was performed to examine the expression patterns of key genes of mesodermal lineages and of early kidney development (Table S1). Relative gene expression levels were evaluated and compared between the following groups: (i) the *Bra*-GFP^+^ and *Bra*-GFP^−^ populations isolated from the EBs (3-D system); (ii) *Bra*-GFP^+^ and *Bra*-GFP^−^ populations isolated from the 2-D system; and (iii) the *Bra*-GFP^+^ populations isolated from the 3-D and 2-D systems. Stemness markers *Oct4* and *Nanog* and the primitive ectoderm marker, *Fgf5*, were also evaluated to assess whether undifferentiated mESCs and/or ectoderm cells were present.

Firstly, comparisons were made between gene expression levels in the *Bra*-GFP^+^ cells and the *Bra*-GFP^−^ cells isolated from the 3-D and 2-D system. The results showed that the early mesoderm genes *Bra*, *Cdx2*, *Tbx6, Foxf1*, *Foxa2*, *Hoxb1* and *Hoxc9* were expressed by *Bra*-GFP^+^ cells isolated from both the 3-D and 2-D systems, but the relative expression levels differed in comparison to the respective *Bra*-GFP^−^ populations. For instance, under the 3-D conditions, the expression levels of *Bra*, *Cdx2*, *Tbx6, Foxf1*, and *Hoxb1* in the *Bra*-GFP^+^ population were approximately 55-, 10-, 40-, 10- and 55-fold higher than in the *Bra*-GFP^−^ population, respectively, whereas under the 2-D conditions, *Bra*, *Tbx6* and *Hoxb1* levels in the *Bra*-GFP^+^ cells were only 2-, 4-, and 5-fold higher, respectively, than in the *Bra*-GFP^−^ cells (Figs 3B–C).

There was a 1- to 10-fold up-regulation of Hox10 and Hox11 paralogy groups (*Hoxa10*, *Hoxa11* and *Hoxd11*) in the *Bra*-GFP^+^ population compared to the *Bra*-GFP^−^ population isolated from cells under 3-D conditions. In contrast, down-regulation of the same genes was observed in the *Bra*-GFP^+^ population isolated from cells under 2-D conditions compared to the *Bra*-GFP^−^ population (Figs 3B–C). This suggested that the status of *Bra*-GFP^+^ cells isolated from EBs may be closer to a stage resembling posterior mesoderm, as it has been shown previously that posterior mesoderm, which gives rise to the MM, expresses higher levels of *Hox10* and *11* genes compared to anterior mesoderm (Taguchi *et al*., 2014).

Genes of intermediate mesoderm and metanephric mesenchyme, i.e., *Lhx1*, *Osr1*, *Pax2* and *Wt1*, displayed a similar trend in the change of relative expression levels between the *Bra*-GFP^+^ and *Bra*-GFP^−^ groups under 3-D and 2-D conditions. It is of note that in the cells isolated from the EBs, *Lhx1* was up-regulated by approximately 10-fold in the *Bra*-GFP^+^ cells compared to the *Bra*-GFP− cells, whereas there was minimal up-regulation in the *Bra*-GFP^+^ cells isolated from the 2-D conditions (Figs 3B–C, Fig. S1). *Oct4, Nanog* and *Fgf5* were also evaluated and the data showed no difference between the *Bra*-GFP^+^ cells and *Bra*-GFP^−^ cells isolated from both 3-D and 2-D conditions (Fig. S1).

Next, the relative expression levels of the various genes in *Bra*-GFP^+^ cells isolated from 3-D and 2-D system was compared. There was no significant difference in the expression levels of *Bra* and *Tbx6*, whereas *Cdx2*, *Foxf1* and *Hoxb1* were significantly up-regulated by 9-, 30-, 5-fold, respectively, in the *Bra*-GFP^+^ cells isolated under 3-D conditions. Another early mesoderm gene *Hoxc9* as well as posterior mesoderm genes *Hox10* and *Hox11* were also up-regulated but not significantly. The expression levels of *Lhx1*, *Osr1*, *Pax2*, *Wt1* and *Gdnf* were comparable between the two populations. On the other hand, *Foxd1*, which, is expressed in MM and stroma, showed a slight 2-fold up-regulation in the 3-D *Bra*-GFP^+^ cells, but this was not statistically significant (Fig. 3D).

### 2.4 *Ex vivo* development of intact and re-aggregated non-chimeric mouse kidney rudiments

In order to evaluate how the *Bra*-GFP^+^ cells behave in the rudiment culture, it was first necessary to establish the typical staining pattern of various renal cell-specific antibodies in intact kidney rudiments cultured *ex vivo*. Following 5 days of *ex vivo* culture, the rudiments were fixed and immunofluorescence was performed to detect the following markers: megalin, which is expressed on the apical surfaces of proximal tubule cells (Ranghini *et al*., 2013; Taguchi *et al*., 2014); Wt1, which is expressed in MM and developing nephrons, and expressed at very high levels in nascent and mature podocytes (Moore *et al*., 1999; Ranghini *et al*., 2013; Taguchi *et al*., 2014); synaptopodin, which is expressed in mature podocytes (Mundel *et al*., 1997; Shankland *et al*., 2007). The rudiments were also stained with rhodamine-labeled peanut agglutinin (PNA), which mainly binds to the basement membranes of UBs, and more weakly to those of the developing nephrons (Laitinen *et al*., 1987). PNA staining showed an intact UB tree, and immunostaining for megalin showed typical staining of the apical surfaces of proximal tubule cells (Fig. 4A). As expected, immunostaining for Wt1 showed weaker expression in MM and developing nephrons and intense expression in nascent and mature podocytes, whereas synaptopodin was exclusively expressed in mature podocytes (Fig. 4A).

**Fig. 4.**
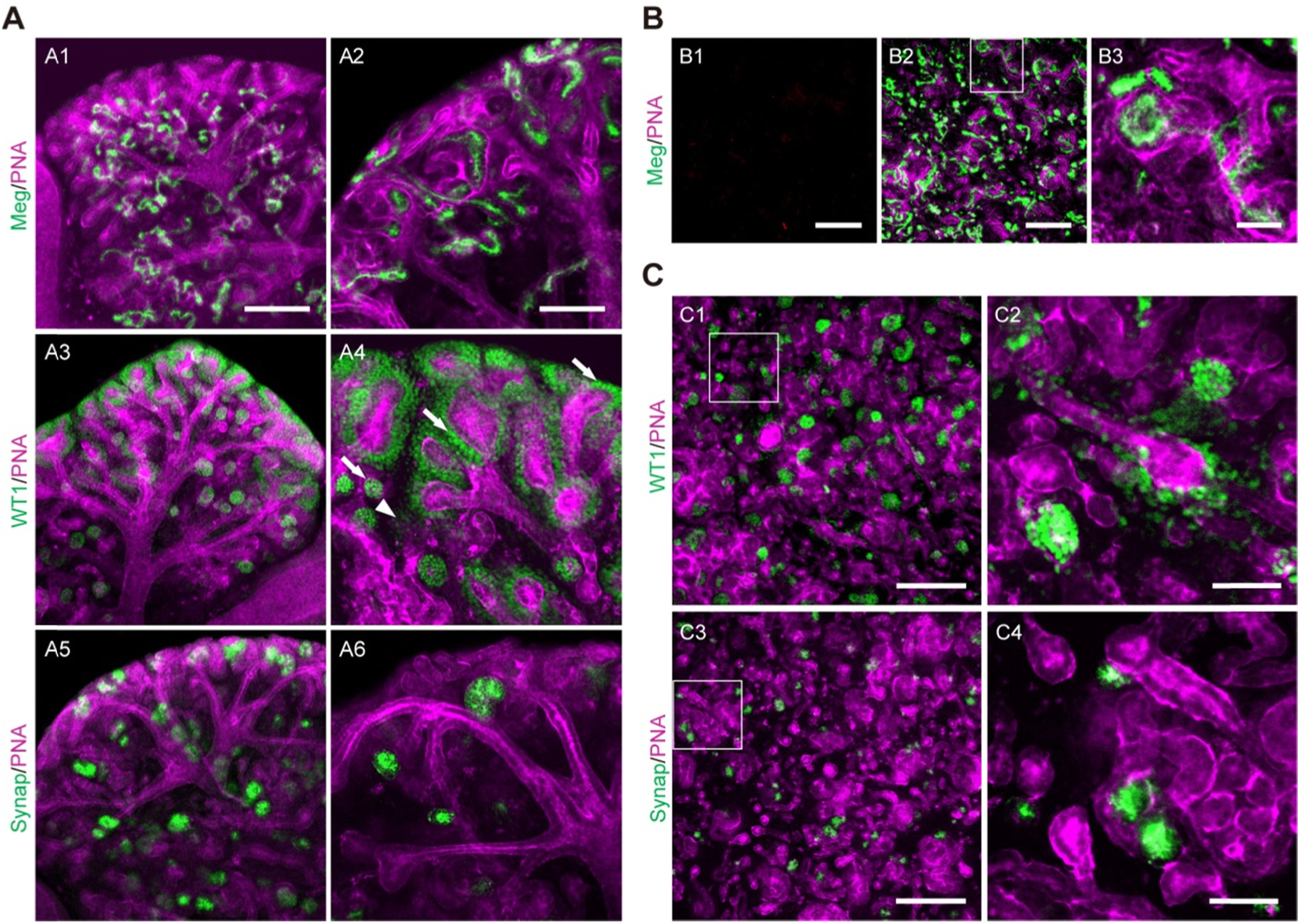
Development of intact E13.5 mouse embryonic kidney and re-aggregated kidney rudiments cultured *ex vivo* for 5 days. (A) Representative confocal photomicrographs of intact kidney showed that proximal tubules were positively stained for megalin (Meg, green) and PNA (magenta). Developing glomeruli were immunostained for Wt1 (green) and synaptopodin (Synap, green) positive staining. Arrows point to developing podocytes and arrowheads point to MM. (B) E13.5 mouse embryonic kidneys were dissociated and pelleted as aggregates comprising 2 ×10^5^ cells for each rudiment. Representative confocal photomicrographs of the re-aggregated rudiments cultured *ex vivo* at days 0 (B1) and 5 (B2-B3) showed that tubule-like structures formed during the 5-day culture. (C) The re-aggregated rudiments contain tubules and nascent glomerular-like structures that are similar to those of the intact rudiments cultured for 5 days. Boxed regions outlined are enlarged in the magnified image. Data were collected from three biological replicates. Scale bars: 200 μm (A1, A3, A5, B1-B2, C1 and C3); 100 μm (A2, A4 and A6); 50 μm (B3, C2 and C4)

To confirm that re-aggregated kidney rudiments could develop nephron and UB structures as previously reported (Unbekandt and Davies, 2010; Rak-Raszewska *et al*., 2012; Ranghini *et al*., 2013), dissociated kidney rudiment cells were pelleted and cultured *ex vivo* prior to staining with the aforementioned markers. Firstly, it was important to confirm that the disaggregation process was effective and that no non-dissociated renal structures were present at the start of the culture period.

Therefore, at day 0, rudiments were stained for megalin and PNA. The results showed that no staining was present at day 0, whereas multiple tubular structures were present by day 5 (Fig. 4B). More detailed analysis of the re-aggregated rudiments showed that the pattern of tubular structures and nascent glomeruli appeared similar to that of the intact rudiments, which was consistent with previous studies (Kuzma-Kuzniarska *et al*., 2012; Rak-Raszewska *et al*., 2012; Ranghini *et al*., 2013). Although UB tubules formed, they did not form a contiguous UB tree (Fig. 4C).

### 2.5 The behaviour of mESC-derived *Bra*-GFP^+^ cells within chimeric kidney rudiments cultured *ex vivo*

Before assessing the differentiation potential of the mESC-derived *Bra*^+^ cells in the chimeric rudiment assay, it was first necessary to confirm that chimeric rudiments comprising a positive control cell population developed as expected. To this end, chimeric rudiments containing GFP^+^ mouse neonatal kidney-derived stem cells (KSCs) were generated, as we have previously shown that KSCs can generate proximal tubule cells and podocytes within rudiments (Ranghini, 2011; Ranghini *et al*., 2013). The chimeric rudiments were cultured for 5 days *ex vivo* and analysed as previously using the renal cell-specific markers. On day 0, the KSCs were evenly distributed in the chimeric rudiments (Fig. S2). After 5 days of culture, the chimeric rudiments had developed proximal tubule-like structures that stained positively for megalin, as well as nascent glomeruli that contained podocytes, as evidenced by positive staining for Wt1 and synaptopodin. KSCs showed integration into the tubules and glomeruli of the developing nephrons (Figs 5–7).

**Fig. 5.**
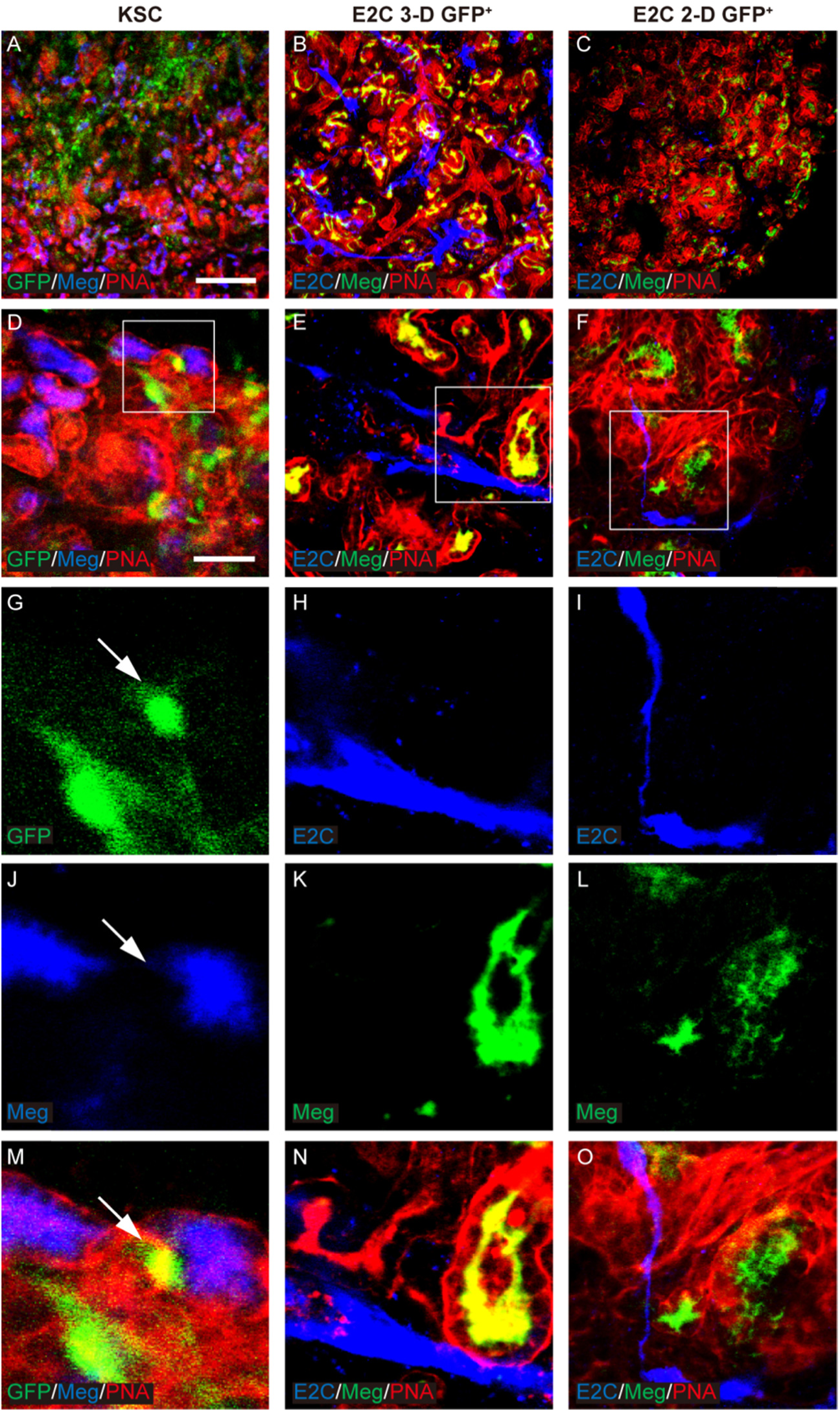
Potential of *Bra*-GFP*/Rosa26-E2C* mESCs isolated from 3-D and 2-D systems to integrate in megalin-expressing renal tubules. Rudiments were cultured *ex vivo* for 5 days. GFP-KSCs (green) were used as positive controls and showed integration into the tubules of the developing nephrons. Arrows point to the GFP^+^ KSCs that had integrated into developing tubules that were dual stained by PNA (red) and megalin (Meg, blue) (G, J and M). In the day-5 chimeric rudiments comprising *Bra*-GFP^+^ cells derived from mESC 3-D system, E2C^+^ *Bra*-GFP^+^ cells (blue) appeared to be elongated and formed an interconnected network within the rudiments. They were often found surrounding the tubules but did not integrate into them (B, E, H, K and N). In the day-5 chimeric rudiments comprising *Bra*-GFP^+^ cells isolated from mESC 2-D system, fewer *Bra*-GFP^+^ cells (blue) were observed, and, unlike those from the 3-D system, most did not appear to be elongated (C, F, I, L and O). Boxed regions outlined are enlarged in the magnified images. Data were collected from three biological replicates. Scale bars, 200 μm (A-C) and 50 μm (D-F).

**Fig. 6.**
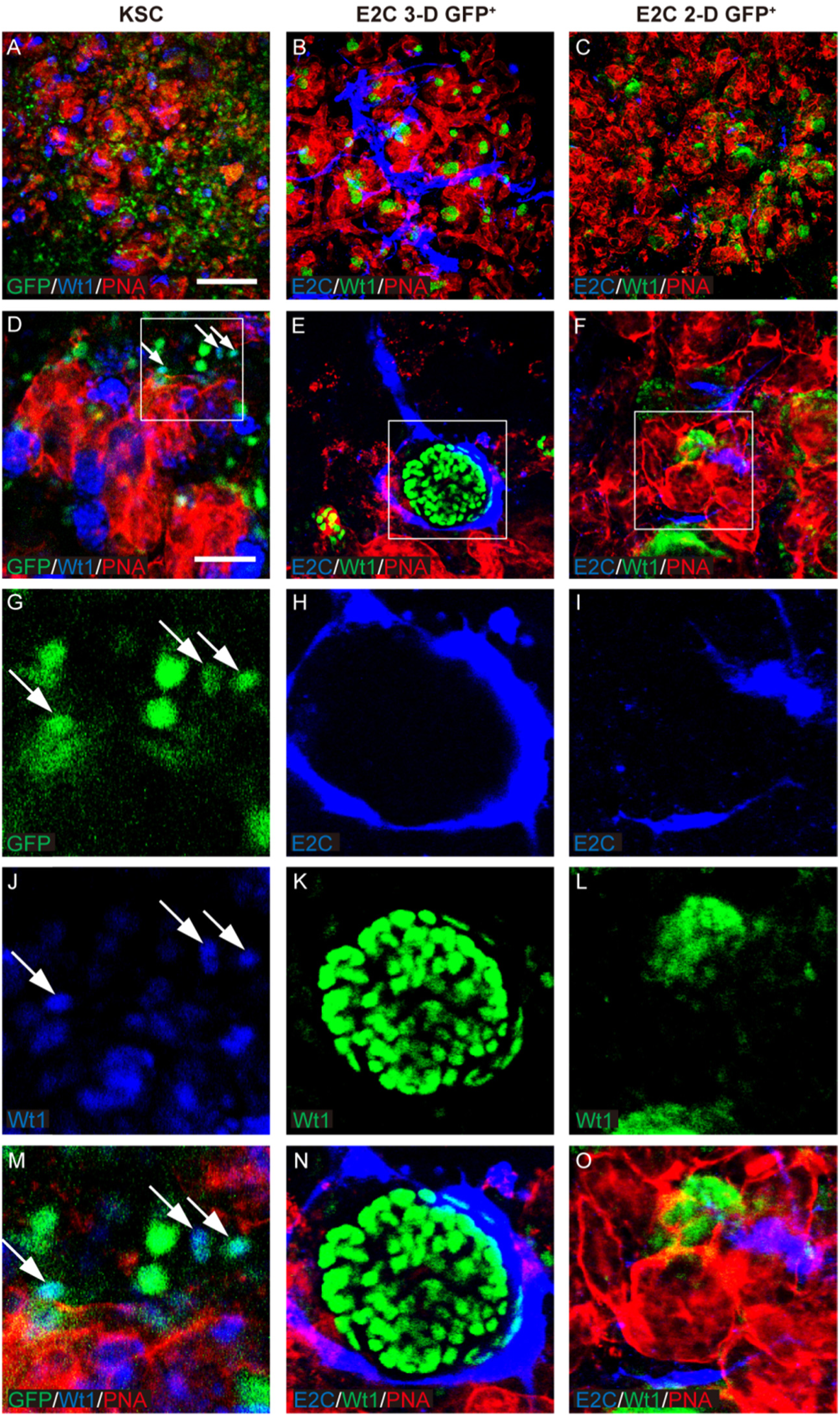
Potential of *Bra*-GFP*/Rosa26-E2C* mESCs isolated from 3-D and 2-D systems to integrate into Wtl-expressing nascent glomeruli. Rudiments were cultured *ex vivo* for 5 days. Arrows point to the integrated KSCs (positive controls) that were GFP-labelled and dual stained by PNA (red) and Wt1 (blue) (D, G, J and, M). E2C^+^ *Bra*-GFP^+^ cells (blue) in the day-5 chimeric rudiments comprising *Bra*-GFP^+^ cells derived from mESC 3-D system were often found surrounding the tubules (red) and glomerular structures (green) but did not integrate into them (B, E, H, K and N). *Bra*-GFP^+^ cells isolated from mESC 2-D system also did not appear to integrate into any renal structures (C, F, I, L and O). Boxed regions outlined are enlarged in the magnified images. Data were collected from three biological replicates. Scale bars, 200 μm (A-C) and 50 μm (D-F).

**Fig. 7.**
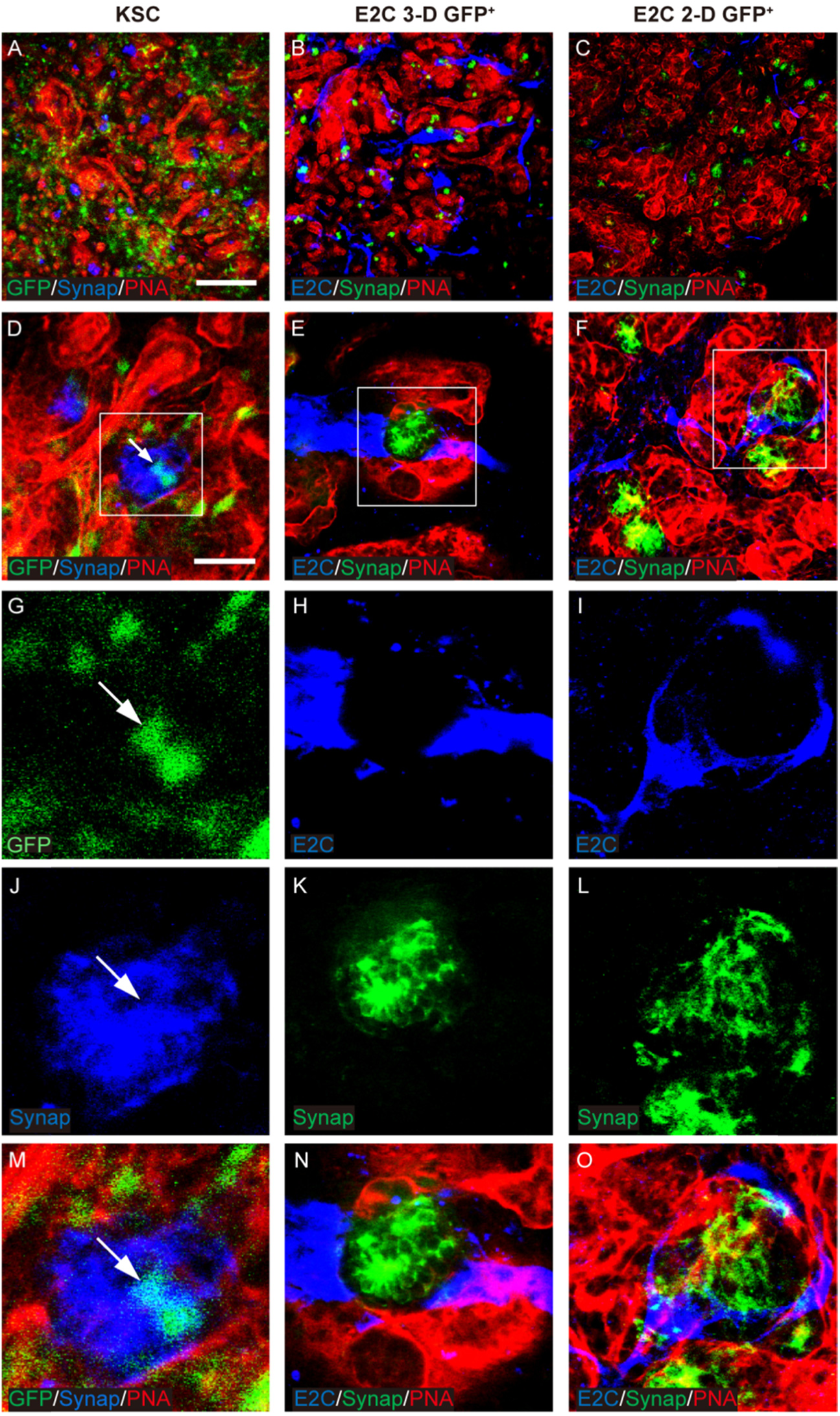
Potential of *Bra*-GFP*/Rosa26-E2C* mESCs isolated from 3-D and 2-D systems to differentiate into synaptopodin-expressing podocytes. Rudiments were cultured *ex vivo* for 5 days. Arrows point to the integrated KSCs (positive controls) that were GFP-labelled and dual stained with PNA (red) and synaptopodin (Synap, blue) (D, G, J and M). In the day-5 chimeric rudiments comprising *Bra*-GFP^+^ cells derived from mESC 3-D system, E2C^+^ *Bra*-GFP^+^ cells (blue) did not generate synaptopodin^+^ cells (B, E, H, K and N). *Bra*-GFP^+^ cells isolated from mESC 2-D system (blue) also failed to generate synaptopodin^+^ cells (C, F, I, L and O). Boxed regions outlined are enlarged in the magnified images. Data were collected from three biological replicates. Scale bars, 200 μm (A-C) and 50 μm (D-F).

To investigate the behaviour of mESC-derived *Bra*-GFP^+^ cells within chimeric kidney rudiments cultured *ex vivo*, Firstly, the behaviour of E2-Crimson-expressing (E2C^+^) *Bra*-GFP^+^ cells isolated from mESC-derived EBs (3-D culture system) were investigated in the *ex vivo* rudiment assay. Staining for PNA, megalin, Wt1 and synaptopodin showed that similarly to the positive control chimeras comprising KSCs, the re-aggregated metanephric cells were able to develop tubular structures and nascent glomeruli (Figs 5–7). However, immunostaining for E2C showed that the EB-derived cells did not integrate into tubules or glomeruli, and instead, appeared to elongate and form interconnected cell networks throughout the rudiment. In many cases, the EB-derived cells appeared to align against the outer surface of developing glomeruli (Figs 5–7).

Next, the behaviour of E2C^+^ *Bra*-GFP^+^ cells isolated from the 2-D culture system was investigated using the chimeric rudiment assay. As with the EB-derived *Bra*-GFP^+^ chimeras, staining for PNA, megalin, Wt1 and synaptopodin showed that re-aggregated metanephric cells in chimeras comprising *Bra*-GFP^+^ cells isolated from the 2-D culture system were able to generate tubular structures and nascent glomeruli (Figs 5–7). Similarly to the E2C^+^ EB-derived *Bra*-GFP^+^ cells, the cells isolated from the 2-D culture system did not appear to integrate into tubules or glomeruli. However, in contrast to the EB-derived cells, those isolated from 2-D culture tended not to form connections with each other. Although elongated cells were occasionally observed in close proximity to developing glomeruli, the majority of the cells were not elongated and did not from interconnected cell networks (Figs 5–7). Furthermore, there appeared to be fewer E2C cells present in these chimeras compared to those generated from mESC-derived *Bra*-GFP^+^ isolated from EBs.

The morphology of E2C^+^ *Bra*-GFP^+^ cells within the chimeras generated from EB-isolated cells appeared similar to that of endothelial cells within *ex vivo* kidney rudiments (Halt *et al*., 2016). To investigate if the E2C^+^ cells had differentiated into endothelial cells, the rudiments were immunostained for the endothelial marker, PECAM-1 (platelet and endothelial cell adhesion molecule 1) (Kondo *et al*., 2007). It was found that the metanephric cells generated PECAM-1^+^ interconnected cell networks in both types of chimeric rudiment, indicating that endothelial cells had differentiated. Analysis of E2C^+^ cells within the chimeric rudiments generated from EB-derived *Bra*-GFP^+^ cells showed that the majority of these cells appeared to stain positively for PECAM-1, suggesting that they had differentiated into endothelial cells. In contrast, most of the E2C^+^ cells within the chimeric rudiments generated from 2-D culture-derived *Bra*-GFP^+^ cells did not stain positively for PECAM-1. Instead, only the elongated cells which were occasionally observed within these chimeras were found to stain for PECAM-1 (Fig. 8, Movies S1–2).

**Fig. 8.**
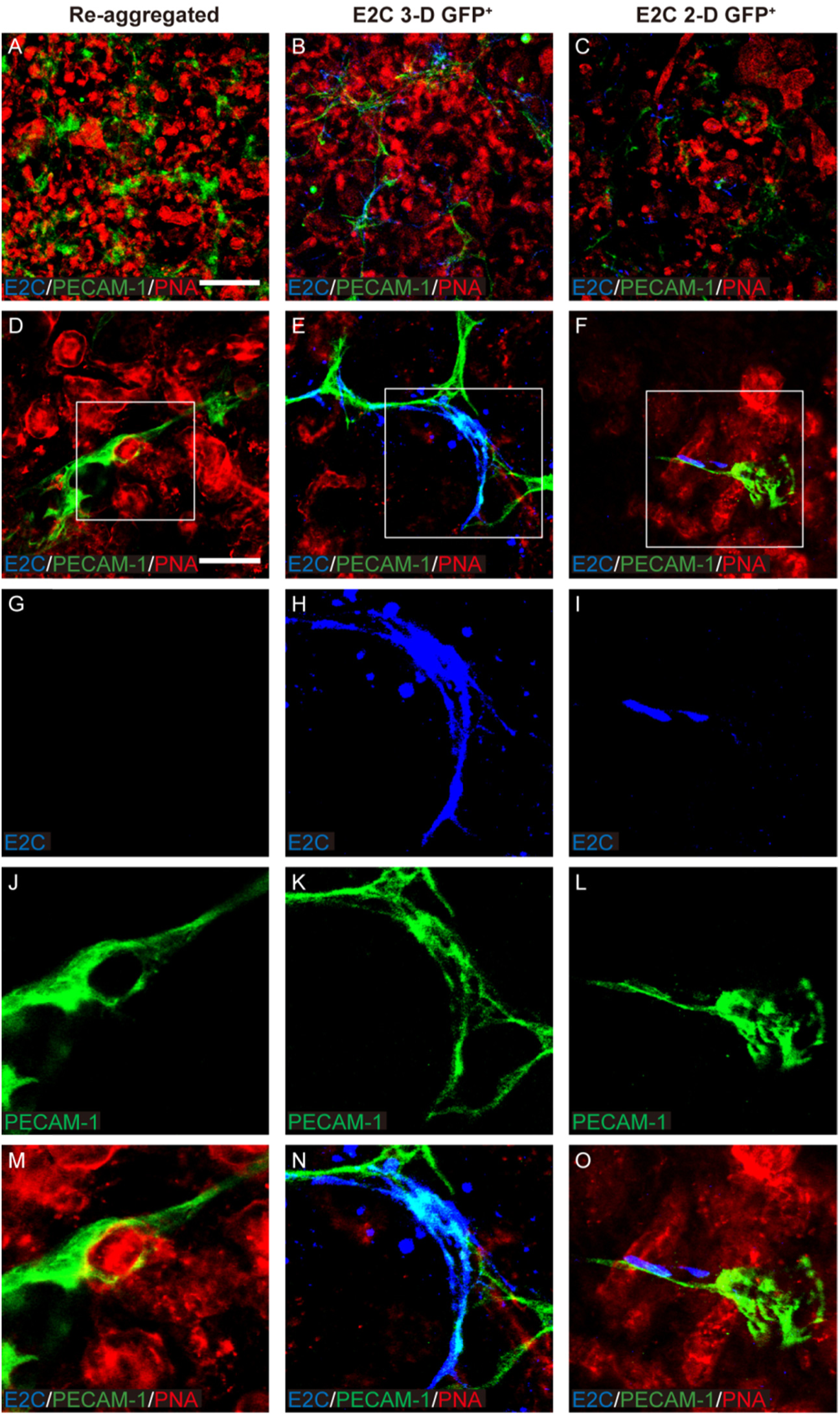
Confocal photomicrographs showing PECAM-1 immunostaining within day-5 *ex vivo* mouse embryonic kidney rudiments comprising *Bra*-GFP^+^ derived from *Bra*-GFP*/Rosa26-E2C* mESCs cultured in 3-D and 2-D systems. Immunostaining for E2C was undertaken to identify the mesodermal cells, and PECAM-1 immunostaining was performed to identify endothelial-like cells. (A, D, G, J, M) Re-aggregated rudiments without exogenous cells; (B, E, H, K, N) Re-aggregated chimeric rudiments containing E2C^+^ *Bra*-GFP^+^ cells isolated from the 3-D culture system; (C, F, I, L, O) Re-aggregated chimeric rudiments containing E2C^+^ *Bra*-GFP^+^ cells isolated from the 2-D culture system. Boxed regions outlined are enlarged in the magnified images. Data were collected from three biological replicates. Scale bars, 200 μm (A-C) and 50 μm (D-F).

## 3. Discussion

In this study, we generated mesoderm populations from a *Bra-GFP/Rosa26-E2C* mESC reporter line using 3-D and 2-D culture systems.

The dynamics of GFP expression during EB culture was similar to what has been previously observed in our group (Rak-Raszewska, 2010); i.e., at low seeding density, GFP appeared to peak earlier than at higher seeding densities. A possible explanation is that mESCs might express inhibitors of mesoderm differentiation, such as noggin, which would be present at higher levels in higher density cultures, and might therefore delay mesoderm differentiation (GFP expression) (Tonegawa and Takahashi, 1998; Gratsch and O’Shea, 2002). Also, GFP expression was detected in EBs generated at low density that had not cavitated. This is similar to our lab’s previous findings using the same *E14-Bra-GFP* mESC line, but with a different culture protocol developed by Fehling et al (Fehling *et al*., 2003; Rak-Raszewska *et al*., 2012). In that study, GFP was only expressed within the EBs during days 3 to 4 with about 60% of the population expressing GFP at day 4 (Rak-Raszewska, 2010). This is much higher than the proportion we observed in the current study (less than 40%). However, EBs generated using Fehling’s method did not form a proamniotic-like cavity, extra-embryonic endoderm or basement membranes. It is therefore envisaged that the properties of *Bra*^+^ mesoderm cells generated from the two types of EBs (i.e., cavitating or non-cavitating), might have different properties and differentiation potential.

An interesting finding from the qRT-PCR analysis was that the expression levels of *Bra* in the GFP^+^ cells isolated from the 3-D system were approximately 50 times higher than in the GFP^−^ cells, but *Bra* levels in GFP^+^ cells isolated from the 2-D system were only approximately three times higher than in the corresponding GFP^−^ cells. Yet despite this, when *Bra* levels in the GFP^+^ cells from the 3-D system were directly compared with levels in GFP^+^ cells from the 2-D system, there was no significant difference. A possible explanation for this is that the GFP^−^ cells in the EBs are likely to be endoderm or ectoderm cells that do not express *Bra*, whereas in the 2-D system, it is possible that the GFP^−^ cells might be committed to the mesodermal lineage and have started to up-regulate *Bra*, but due to the time-lag between transcription and translation, might not have yet started to produce GFP. If this were the case, such cells would be *Bra*^+^ but GFP^−^, and would thus have been sorted into the GFP-negative fraction by FACS.

When comparing the expression levels of key genes between the GFP^+^ cells from the 3-D and 2-D systems, there were only three genes that were significantly up-regulated in the cells from 3-D system, namely, *Foxf1*, *Cdx2* and *Hoxb1*. The high expression levels of *Foxf1* might suggest that the GFP^+^ cells from the 3-D system might be lateral plate mesoderm cells. It is known that high levels of BMPs promote the differentiation of lateral plate mesoderm, whereas low levels of BMPs promote intermediate mesoderm (Tonegawa and Takahashi, 1998). It is therefore possible that in the larger cavitating EBs, there might be higher levels of BMPs which would then drive the differentiation of lateral plate mesoderm. However, the cells also had significantly higher levels of the nascent mesoderm gene, *Cdx2*, and the posterior mesoderm gene, *Hoxb1*. Furthermore, although not significant, there was a clear trend that the *Hox* genes tested, which are expressed in intermediate mesoderm, were up-regulated in the cells from the 3-D system.

By introducing the E2C-expressing mesodermal cells into the chimeric rudiments *ex vivo*, we showed that neither the *Bra*-GFP^+^ cells derived from the 3-D nor 2-D culture systems appeared to integrate into the developing nephrons. The results are strikingly different from our lab’s previous studies that investigated the nephrogenic potential of *Bra*-GFP^+^ cells isolated from non-cavitating EBs in the same rudiment culture assay (Rak-Raszewska *et al*., 2012). In these earlier studies, it was found that *Bra*-GFP^+^ mESCs derived from non-cavitating EBs were able to integrate into both the developing nephrons and UBs, and could form functional proximal tubule cells and podocytes (Rak-Raszewska *et al*., 2012). Another study by Vigneau *et al* showed that *Bra*^+^ cells derived from mouse EBs contributed to the proximal tubules when injected into the neonatal mouse kidney *in vivo* (Vigneau *et al*., 2007). The results we obtained with the *Bra*-GFP^+^ cells obtained from cavitating EBs were surprising. We had expected that as these cells were isolated at a later time point than the *Bra*-GFP^+^ cells in the non-cavitating EBs, they might more closely resemble posterior mesoderm, which has recently been shown to generate the MM but not the UB (Taguchi *et al*., 2014). We therefore thought that these cells might integrate into developing nephrons, but not the UBs. However, they did not integrate into either of these structures and instead appeared to differentiate into endothelial cells. There have been contrasting reports concerning the presence of endothelial cells in mouse kidney rudiments cultured *ex vivo*, with some studies suggesting endothelial cells cannot survive in *ex vivo* rudiments (Loughna *et al*., 1997) and others suggesting they do (Halt *et al*., 2016). Our findings are consistent with the Halt *et al* study that indicates endothelial cells are present in rudiments, and similarly to that study, we found that although the endothelial cells formed interconnected networks, they did not form capillaries with lumen, nor did they invest the developing glomeruli.

The key differences in the gene expression profile of the *Bra*-GFP^+^ cells isolated from cavitating EBs (current study) and non-cavitating EBs (previous study) (Rak-Raszewska, 2010) is that in comparison to GFP^−^ cells, the former expressed much higher levels of *Foxf1*, which is highly expressed in lateral plate mesoderm, and lower levels of the MM genes, *Gdnf* and *Osr1* (Rak-Raszewska, 2010). The high expression levels of *Foxf1* might explain why the EB-derived *Bra*-GFP^+^ cells in the current study had a tendency to generate endothelial cells, because it is known that *Foxf1* is essential for vasculogenesis in the developing embryo and is expressed in endothelial cells (Mahlapuu *et al*., 2001; Ren *et al*., 2014).

High levels of BMP signals and their receptors ALK3/6 have been shown to promote a lateral plate mesoderm fate (James and Schultheiss, 2005). Due to the heterogeneous nature of the EBs, it is possible that mesoderm niches that resemble dynamic microenvironments of the *in vivo* primitive streak have been formed. Cells residing in the niches that are exposed to high concentrations of BMP signals might, therefore, adopt a lateral plate mesoderm fate. Retinoic acid, FGF and Wnt signals might also affect the cell commitment of lateral plate mesoderm but their effects may be stochastic within the EBs. Nevertheless, we cannot exclude the possibility that the timing might have been another factor; for instance, *Bra*-GFP^+^ cells isolated at slightly earlier or later time-points might have expressed genes of other mesodermal lineages.

Regarding the *Bra*-GFP^+^ isolated from the 2-D system, it was found that these also did not integrate into developing nephrons or UBs. Furthermore, only a small proportion of these cells appeared to differentiate into endothelial cells. The majority of the cells did not form interconnected cell networks and appeared to be randomly dispersed throughout the stroma. Similarly to the *Bra*-GFP^+^ cells from the cavitating EBs, the *Bra*-GFP^+^ cells from the 2-D system did not show any noticeable up-regulation of *Gdnf* or *Osr1* in comparison with the *Bra*-GFP^−^ cells. However, in contrast to the EB-derived cells, those isolated from the 2-D system did not show up-regulation of *Foxf1*, which is consistent with their limited tendency to generate endothelial cells. It is possible that the *Bra*-GFP^+^ cells from the 2-D system might have differentiated into stromal cells, but it was not possible to test this due to the lack of a stroma-specific antibody. It is interesting to note that the *Bra*-GFP^+^ cells from the 2-D system expressed higher levels of the stromal gene, *Foxd1* (Mugford *et al*., 2008) compared to those from the 3-D system, but the results were not statistically significant.

## 4. Materials and methods

### 4.1 Routine cell culture

*Bra-GFP/Rosa26-E2C* mESCs (Zhou *et al*., in press) were maintained in 0.1% gelatinised 6-well tissue culture plates with mitomycin-C (Sigma-Aldrich, M4287) inactivated STO (ATCC, SCRC-1049) feeder cells at 37°C in a humidified incubator with 5% CO_2_ in Dulbecco’s Modified Eagle’s Medium (DMEM) (Sigma-Aldrich, D6546) supplemented with 150 mL L^−1^ FBS (Sigma-Aldrich, F2442), 10 mL L^−1^ MEM non-essential amino acid (Sigma-Aldrich, M7145), 10 mL L^−1^ L-glutamine (Sigma-Aldrich, G7513), 0.1 mmol L^−1^ β-mercaptoethanol (Gibco, 31350) and 1,000 U mL^−1^ mouse leukemia inhibitory factor (mLIF) (Merck Millipore, ESG1107). Cells were passaged every other day and those at passage 13–22 were used for experiments.

GFP-expressing mouse neonatal kidney-derived stem cells (GFP-KSCs) (Ranghini, 2011) were maintained in 60 mm tissue culture dishes at 37°C in a humidified incubator with 5% CO_2_ in DMEM supplemented with 100 mL L^−1^ FBS (Gibco, 10270), 10 mL L^−1^ MEM non-essential amino acid (Sigma-Aldrich, M7145), 10 mL L^−1^ L-glutamine (Sigma-Aldrich, G7513) and 0.1 mmol L^−1^ β-mercaptoethanol (Gibco, 31350). Cells were passaged 2–3 times per week and those at passage 17– 20 were used for experiments.

### 4.2 3-D EB system

mESCs were sub-cultured in gelatinised 6-well tissue culture plates for 48 h to deplete feeder cells. Cells were then collected and plated in 90 mm bacterial petri dishes (Sterilin, 101VR20) at the densities of 6.25×10^4^, 1.25×10^5^ and 2.5×10^5^ cells mL^−1^ to form aggregates. The EBs were maintained at 37°C in a humidified incubator with 5% CO_2_ in DMEM supplemented with 100 mL L^−1^ FBS (Sigma-Aldrich, F2442), 10 mL L^−1^ MEM non-essential amino acid, 10 mL L^−1^ L-glutamine and 0.1 mmol L^−1^ β-mercaptoethanol for up to 9 days with a medium change every other day. Each dish was split 1:2 on day 3and EB morphology was examined on days 4 and 7. Experiments were carried out in 3 independent biological replicates.

### 4.3 2-D system

mESCs were sub-cultured in gelatinised 6-well tissue culture plates for 48 h to deplete feeder cells. Cells were collected and plated into gelatinised 6-well plates at 1×10^5^ cells per cm^2^ for 24 h. 2-D induction culture was based on the protocols previously described (Turner *et al*., 2014a,b). Briefly, cells were then harvested and re-plated into 60 mm tissue culture dishes at a density of 4.7×10^3^ cells per cm^2^ with overnight incubation in mESC culture medium. The following morning, medium was changed to NDiff^®^ 227 (Clontech, Y40002) for 48 h and then to NDiff^®^ 227 supplemented with Activin-A (R&D Systems, 338-AC) and CHIR 99021 (Tocris, 4423) to a final concentration of 100 ng mL^−1^ and 3 µmol L^−1^, respectively, for a further 48 h incubation. Medium was changed on a daily basis. Experiments were carried out in 3 independent biological replicates.

### 4.4 Cell-IQ real-time imaging

On day 3, EBs that were formed from mESCs at the plating density of 1.25×10^5^ cells mL^−1^ were harvested and plated onto solidified 2% agarose gel (Sigma-Aldrich, A9045) in glass bottom 6-well plates (MatTek, P06G-0–20-F). They were then embedded in a thin overlay of 1% agarose. Each well was filled with 3 mL EB medium once the overlaid gels were set. Plates were maintained in Cell-IQ (Chip-Man Technologies Ltd) imaging facility. EBs were imaged by the Cell-IQ Imagen (Chip-Man Technologies Ltd) software on days 3 to 9 on an hourly basis. Imaging data from both bright field and 488 nm laser for the GFP fluorescence signal were documented from 3 independent biological replicates. Raw data were analysed by the Cell-IQ Analyser (Chip-Man Technologies Ltd) and ImageJ (NIH) softwares.

### 4.5 EB fixation and cryo-sectioning

EBs were harvested on day 7 and fixed with 4% paraformaldehyde (PFA). They were then soaked in 15% sucrose followed by embedding in the 7.5% molten gelatin. Samples were mounted onto cork disks with Shandon™ Cryomatrix™ embedding resin (Thermo Fisher Scientific, 6769006) and cut with a cryostat at 20 µm.

### 4.6 Flow cytometry analysis

Single cell suspensions of 1×10^6^ cells mL^−1^ were obtained from 3-D or 2-D culture systems and examined by a BD FACScalibur (BD Biosciences) flow cytometer according to manufacturer’s instructions, using a 488 nm laser to detect the GFP signal. For analysis of the GFP expression window in the EBs, wild-type E14TG2a-derived EBs were used as a negative control. For analysis of GFP expression in the 2-D system, undifferentiated *Bra-GFP/Rosa26-E2C* mESCs sub-cultured in gelatinised dishes in mESC medium for 24 h prior to induction were used as a negative control. Data were acquired from two biological replicates by the BD CellQuest (BD Biosciences) software based on 10^4^ events and analysed using the Cyflogic (CyFlo Ltd, version 1.2.1) software.

### 4.7 Fluorescence-activated cell sorting (FACS)

Single cell suspensions of 1×10^7^ cells mL^−1^ were obtained from day-6 3-D EBs or day-4 2-D monolayer cultures. Sorting was performed to isolate *Bra*-GFP^+^ cells using the BD FACSAria (BD Biosciences) flow sorter with the 530/30 bandpass filter and 502 longpass mirror. Day-6 EBs derived from wild-type E14TG2a mESCs and undifferentiated *Bra-GFP/Rosa26-E2C* mESCs sub-cultured in gelatinised dishes for 24 h prior to induced differentiation were used as negative controls for 3-D and 2-D systems, respectively. Data output was performed using BD FACSDiva (version 6.1.3) software. Experiments were performed in 3 independent biological replicates.

### 4.8 qRT-PCR and statistical analysis

Cell lysis of FACS-sorted *Bra*-GFP^+^ populations, reverse transcription and qPCR amplification was performed using the Fast SYBR^®^ Green Cells-to-CT™ Kit (Thermo Fisher Scientific, 4405659) in accordance with the manufacturer’s instructions. Gene transcription was detected by the Bio-Rad CFX Connect Real-time PCR Detection System (Bio-Rad) using specific primers validated in-house (Table S2). The reaction was set up with the following steps: 95°C for 20 s initial DNA polymerase activation followed by 40 cycles of denaturation at 95°C for 3 s and annealing/extension at 60°C for 30 s. qPCR specificity was assessed by melt curves and then verified by agarose gel electrophoresis. Non-template control was performed for each analysed gene and the non-reverse transcriptase control was also included to verify the elimination of genomic DNA. Three biological replicates for the *Bra*-GFP^+^ populations isolated from 3-D and 2-D systems, and two biological replicates for *Bra*-GFP^−^ populations derived from the 3-D and 2-D systems were assessed. For each reaction product analysed, two technical replicates were prepared. Data were acquired using the incorporated Bio-Rad CFX Manager (version 3.1) software. Relative gene expression levels normalised to two endogenous reference genes *Gapdh* and *β-actin* (ΔΔCt) and statistical analysis were also performed using two-tailed Student's *t*-test by the same software, where *P*<0.05 was considered statistically significant.

### 4.9 Mouse embryonic kidney rudiment *ex vivo* culture

The Mouse embryonic kidney rudiment *ex vivo* culture was based on the protocols previously described (Unbekandt and Davies, 2010). Briefly, kidneys were dissected out from embryonic day (E) 13.5 CD1 mouse (Charles River) and dissociated into single cells following an incubation of 15 min in 0.25% trypsin/PBS (Sigma-Aldrich, T4174) with intermittent gentle agitation. Cells were pelleted at 1 800 ×g for 2 min and re-suspended in kidney rudiment medium comprising MEME (Sigma-Aldrich, M5650) and 100 mL L^−1^ FBS. In the meantime, FACS-sorted *Bra*-GFP^+^ cells derived from mESC 3-D or 2-D systems were collected in rudiment medium and counted. A total of 2×10^5^ cells were used in each rudiment, wherein kidney rudiment cells and *Bra*-GFP^+^ cells were mixed at a ratio of 1:9. Rudiments were cultured with Rho-associated, coiled-coil containing protein kinase inhibitor (ROCKi, Y-27632, Merck Millipore, 688001) for 24 h followed by a further 4-day in the absence of ROCKi. Controls were also set up, including kidney rudiments comprising GFP-KSCs (1:9 ratio of KSC: kidney rudiment cells), reaggregated kidney rudiments (formed by kidney rudiment cells only), and intact kidney rudiments. Experiments were performed in 3 independent biological replicates.

### 4.10 Immunofluorescence staining

For EB frozen section assay, sections were blocked in 10% serum solution and incubated with E2C primary and secondary antibodies followed by nuclear counter-staining of 4’,6-diamidino-2-phenylindole (DAPI, Thermo Fisher Scientific, D1306, 1/100 000). Slides were mounted with DAKO fluorescent mounting medium (Agilent Technologies, S3023) and sealed for viewing on the Leica DM2500 (Leica) fluorescence microscope with a 40× objective and appropriate excitation and emission filter sets. Data were acquired using the Leica Application Suite (LAS, Leica) integrated software and analysed by the ImageJ (NIH, version 1.50i) software.

For mouse embryonic kidney rudiments assay, immunofluorescence and image analysis were carried out based on the protocols described previously (Rak-Raszewska *et al*., 2012; Ranghini *et al*., 2013). Briefly, rudiments of days 0 and 5 were fixed with 4% PFA and blocked with 10% serum solution containing 0.1% Triton-X 100, followed by incubation with primary antibodies for E2C, megalin, Wt1, synaptopodin and PECAM-1, where necessary. They were then incubated with secondary antibodies followed by counter-staining of 10 µg µL^−1^ PNA (Vector, RL–1072). Controls were also included as above to check for non-specific binding of secondary antibodies. Samples were mounted with DAKO fluorescent mounting medium (Agilent Technologies, S3023) and sealed. Data were acquired using the Zeiss LSM 510 META (Zeiss) multiphoton confocal laser scanning microscope with a 40× oil immersion, 20× or 10× lens and appropriate excitation and emission filter sets. Image data analysis was performed by the ImageJ (NIH) and Imaris (Bitplane, version 9.0.2) softwares.

The following primary antibodies were used: rabbit polyclonal IgG E2C (Clontech, 632496, 1/1 000), mouse monoclonal megalin IgG1 (Acris, DM3613P, 1/200), mouse monoclonal Wt1 (Millipore, 05– 753, 1/100), mouse monoclonal synaptopodin IgG_1_ (Progen, 65194, 1/2), rat monoclonal PECAM (BD Pharmingen, 550274, 1/100). Secondary antibodies used were: Alexa Fluor (AF) 488-conjugated chicken anti-rabbit IgG (Thermo Fisher Scientific, AF A21441, 1/1 000), AF594 goat anti-rabbit (Thermo Fisher Scientific, AF A11012, 1/1 000), AF488 goat anti-mouse IgG_1_ (Thermo Fisher Scientific, AF A21121, 1/1 000), AF647 donkey anti-mouse IgG (H+L) (Thermo Fisher Scientific, AF A31571, 1/1 000), AF488 donkey anti-rat IgG (H+L) (Thermo Fisher Scientific, AF A21208, 1/ 1 000).

## Acknowledgements

The authors thank Dr. Sandra Pereira Cachinho, for her help with the FACS, Dr. Marco Marcello and Dr. Joanna Wnetrzak, for their assistance on the confocal laser scanning microscopy, and Dr. David Mason, for his advice on 3-D microscopy image processing, respectively. The authors also acknowledge the support by the Biomedical Services Unit as well as the Cell Sorting and Isolation Facilities and the Centre for Cell Imaging of the Technology Directorate at the University of Liverpool.

## Supplementary Figures

**Fig. S1.**
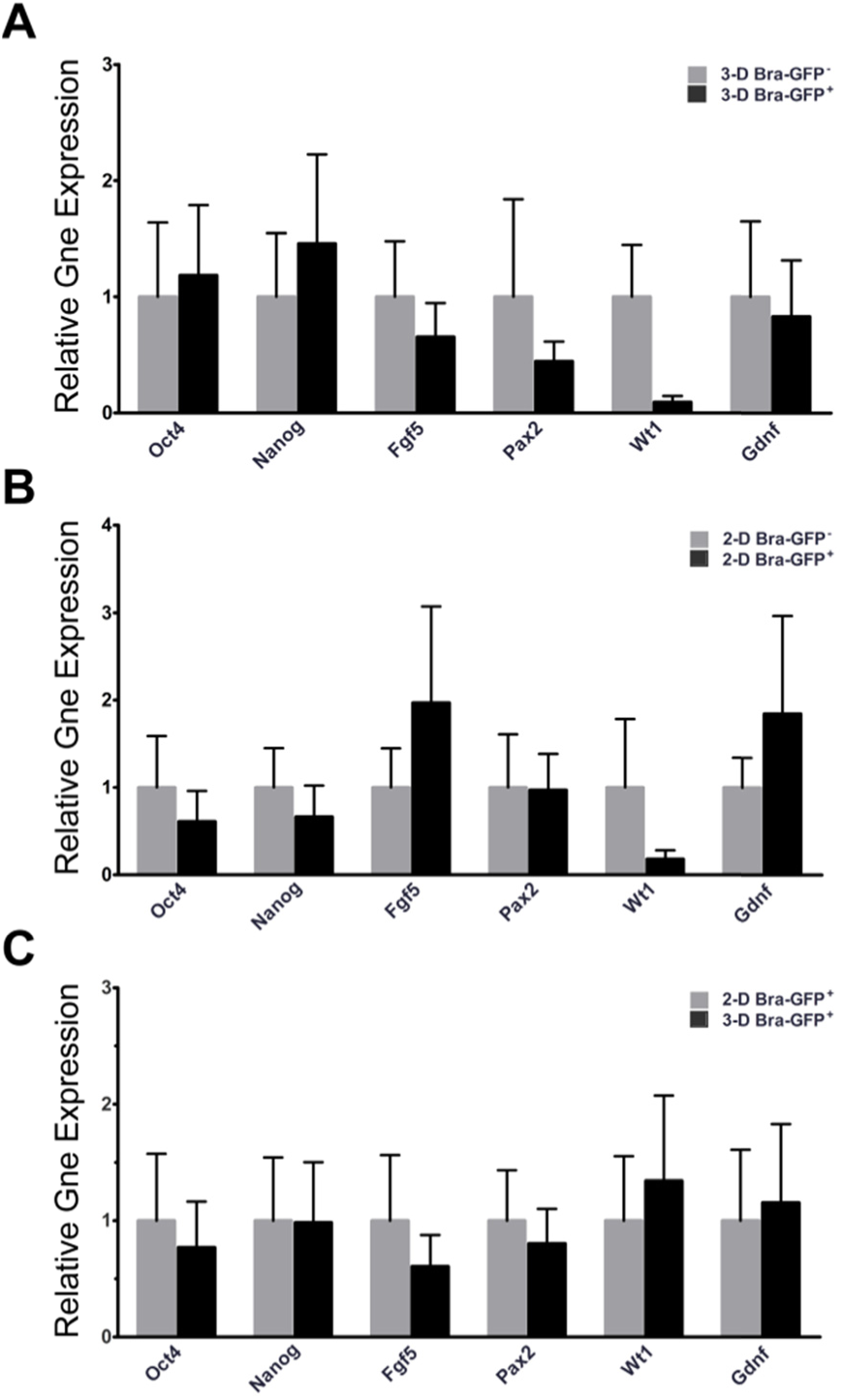
qRT-PCR analysis of stemness and lineage markers expressed by the mesodermal and non-mesodermal populations from *Bra*-GFP*/Rosa26-E2C* mESCs cultured in 3-D and 2-D systems. (A) Relative expression levels of genes were compared between *Bra*-GFP^+^ and *Bra*-GFP^−^ populations isolated from the 3-D system (n=2 biological replicates), presented as mean±s.e.m. Data were not statistically assessed on significance due to 2 biological replicates however they gave an indication of the difference between *Bra*-GFP^+^ and *Bra*-GFP^−^ populations. (B) Relative expression levels of genes were compared between *Bra*-GFP^+^ and *Bra*-GFP^−^ populations isolated from the 2-D system (n=2 biological replicates), presented as mean±s.e.m. Data were not statistically assessed on significance due to 2 biological replicates however they gave an indication of the difference between *Bra*-GFP^+^ and *Bra*-GFP^−^ populations. (C) Relative gene expression levels genes were compared between *Bra*-GFP^+^ populations isolated from 3-D system (n=3 biological replicates) and 2-D system (n=3 biological replicates), presented as mean±s.e.m. *P*<0.05 was considered as statistically significant (*t*-test). No significant difference was found between the two systems.

**Fig. S2.**
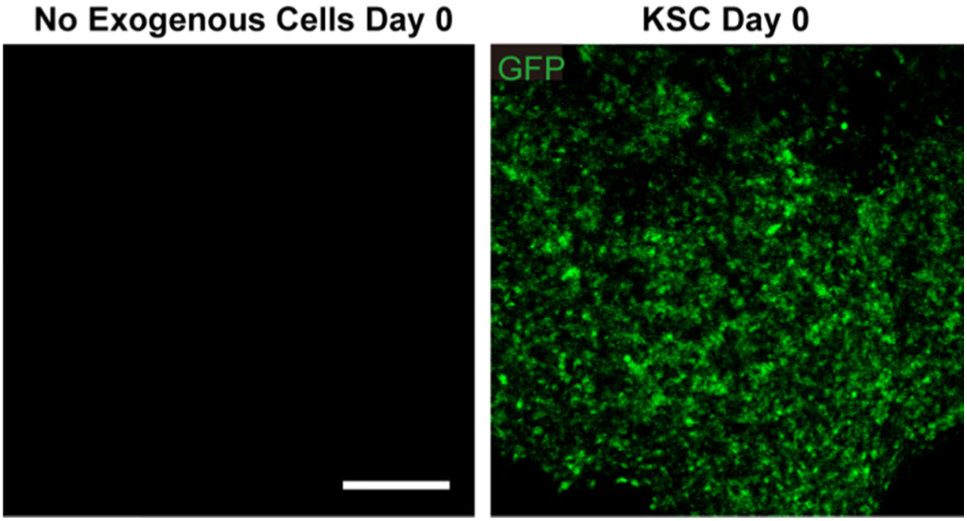
Confocal photomicrographs of the re-aggregated E13.5 mouse embryonic kidney rudiments at day 0 of *ex vivo* culture containing no exogenous cells or GFP-KSCs. GFP-KSCs (positive controls) showed even distribution represented by GFP (green) in the rudiments at the beginning of the culture. Data were collected from three biological replicates. Scale bar, 200 μm

## Supplementary Tables

**Table S1.** List of key genes investigated by qRT-PCR in this study

| Genes | Expression Regions | References | Genes | Expression Regions | References |
| --- | --- | --- | --- | --- | --- |
| *Bra* | PS, TB, notocord | a, b | *Foxa2* | Anterior PS | m |
| *Tbx6* | PS, PM, TB | a-c | *Foxd1* | MM stroma | n |
| *Cdx2* | PS | d-f | *Foxf1* | LPM | o |
| *Lhx1* | LPM, IM | g | *Hoxa10* | PM, MM | e, p, q |
| *Osr1* | LPM, IM, MM | e, g | *Hoxa11* | PM, MM | e, p, q |
| *Pax2* | IM, ND, MM | g, h | *Hoxb1* | Posterior PS | m, r |
| *Wt1* | IM, MM | i | *Hoxc9* | Posterior PM | s |
| *Gdnf* | MM | j-l | *Hoxd11* | PM, MM | p, q |

Notes: PS, primitive streak; PM, paraxial mesoderm; LPM, lateral plate mesoderm; IM, intermediate mesoderm; ND, nephric duct; MM, metanephric mesenchyme; TB, tailbud.
References: a, Papaioannou, 2014; b, Herrmann *et al.*, 1990; c, Chapman *et al.*, 2003; d, Arnold and Robertson, 2009; e, Taguchi *et al.*, 2014; f, Savory, *et al.*, 2009; h, James *et al.*, 2005; i, Little, 2015; j, Lin *et al.*, 1993; k, Sanchez *et al.*, 1996; l, Basson *et al.*, 2006; m, Gadue *et al.*, 2006; n, Mugford *et al.*, 2008; o, Mahlapuu *et al.*, 2001; p, Carapuço *et al.*, 2005; q, Yallowitz *et al.*, 2011; r, Kmita *et al.*, 2000; s, Erselius *et al.*, 1990.

**Table S2.**
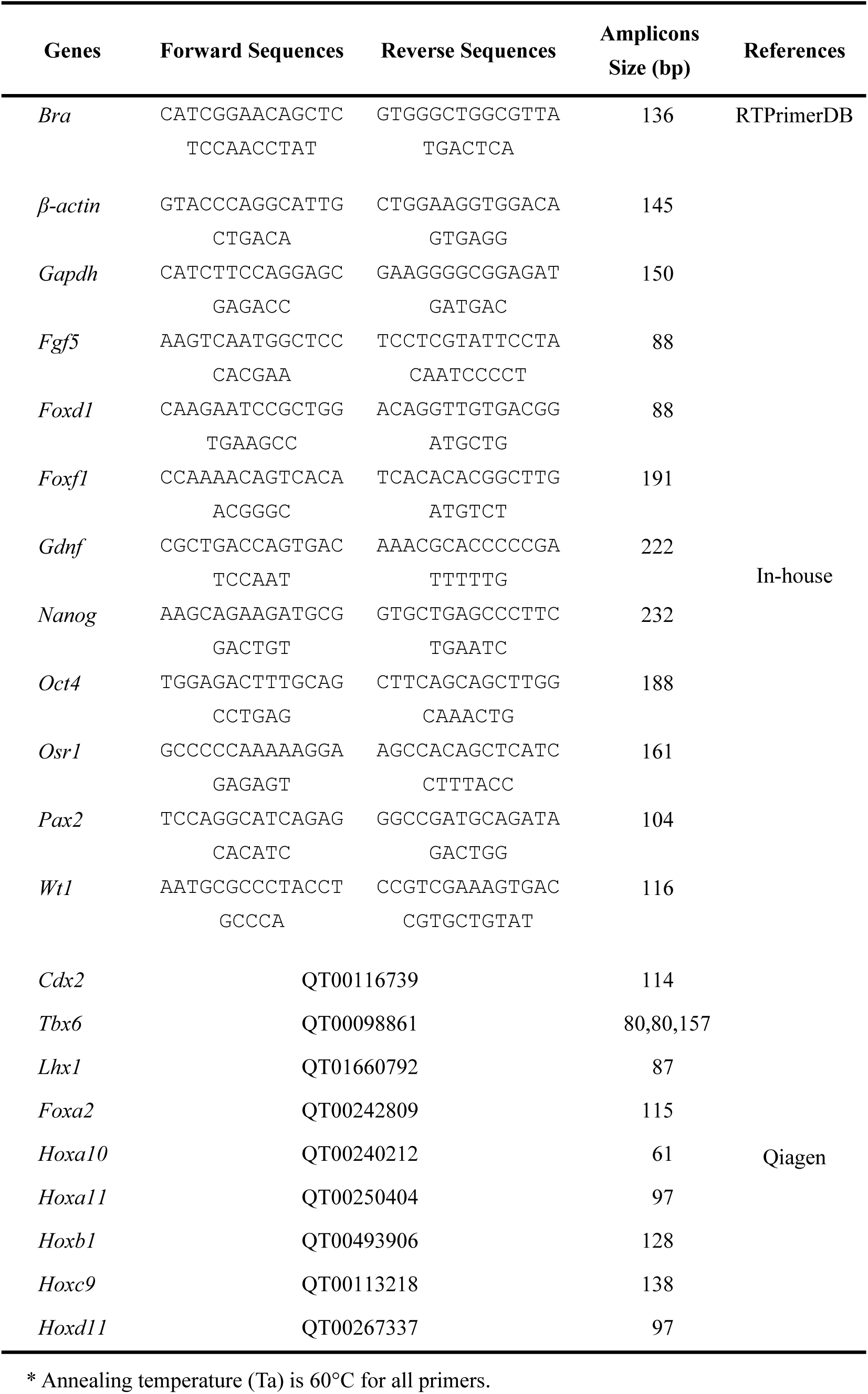
List of qRT-PCR primers

## Supplementary Movies

**Movie S1: Representative 360-degree horizontal 3-D construction of confocal photomicrographs showing spatial distribution of PECAM-expressing 3-D system-derived E2C^+^ *Bra*-GFP^+^ cells within mouse embryonic kidney rudiments.** Rudiments were cultured *ex vivo* for 5 days. Immunostaining for E2C (blue) and PECAM-1 (green) was performed to identify mesodermal and endothelial-like cells, respectively (attached as a separate file).

**Movie S2: Representative 360-degree horizontal 3-D construction of confocal photomicrographs showing spatial distribution of PECAM-expressing 2-D system-derived E2C^+^ *Bra*-GFP^+^ cells within mouse embryonic kidney rudiments.** Rudiments were cultured *ex vivo* for 5 days. Immunostaining for E2C (blue) and PECAM-1 (green) was performed to identify mesodermal and endothelial-like cells, respectively (attached as a separate file).

